# Macrophage-mediated antibody dependent effector function in aggressive B-cell lymphoma treatment is enhanced by Ibrutinib via inhibition of JAK2

**DOI:** 10.1101/2020.06.10.135632

**Authors:** Verena Barbarino, Sinika Henschke, Stuart James Blakemore, Elena Izquierdo, Michael Michalik, Nadine Nickel, Indra Möllenkotte, Daniela Vorholt, Reinhild Brinker, Oleg Fedorchenko, Nelly Mikhael, Tamina Seeger-Nukpezah, Michael Hallek, Christian P. Pallasch

## Abstract

Targeted inhibition of Bruton’s Tyrosine Kinase (BTK) with ibrutinib and other agents has become important treatment options in chronic lymphocytic leukemia, Waldenström’s Macroglobulinemia, Mantle cell lymphoma and non-GCB DLBCL. Clinical trials combining small molecule inhibitors with monoclonal antibodies have been initiated at rapid pace, with the biological understanding between their synergistic interactions lagging behind. Here, we have evaluated the synergy between BTK inhibitors and monoclonal antibody therapy via macrophage mediated antibody dependent cellular phagocytosis (ADCP). Initially, we observed increased ADCP with ibrutinib, whilst second generation BTK inhibitors failed to synergistically interact with monoclonal antibody treatment. Kinase activity profiling under BTK inhibition identified significant loss of Janus Kinase 2 (JAK2) only under ibrutinib treatment. We validated this potential off-target effect via JAK inhibition *in vitro* as well as with CRISPR/Cas9 JAK2^−/−^ experiments *in vivo*, showing increased ADCP and prolonged survival, respectively. This data supports inhibition of the JAK-STAT signaling pathway in B-cell malignancies in combination with monoclonal antibody therapy to increase macrophage mediated immune responses.

## Introduction

Inhibition of Bruton’s Tyrosine Kinase (BTK) in the treatment of B-cell malignancies has been an exemplary story of functional understanding of disease pathogenesis translating to superior survival rates (1–4). Moreover, inhibition of BTK is currently also being evaluated as a therapeutic strategy for patients with severe COVID-19 disease (5,6). BTK is localized in proximity of the B-cell receptor (BCR), forming a signalosome complex upon BCR activation with other BCR associated kinases (BAK); LYN, SYK, and PI3K (7,8). The first generation BTK inhibitor (BTKi) ibrutinib covalently binds its target, which leads to prolonged lymphocytosis via reduced malignant B-cell homing capacity and chemokine signaling (9). However, ibrutinib also has high affinity to other Tec kinases, leading to the development of second generation BTKis, such as tirabrutinib and acalabrutinib, which are highly selective for BTK (10,11).

After initial success of ibrutinib in early phase clinical trials as a single agent in the relapse/refractory setting, clinical trials were quickly initiated in combination with other frontline agents, namely monoclonal antibodies targeting CD20 (anti-CD20; such as rituximab) (12–15). The most recent phase 3 trial in previously untreated chronic lymphocytic leukemia (CLL) offers superior survival in comparison to the chemo-immunotherapeutic (CIT) regimen fludarabine cyclophosphamide and rituximab (FCR) (12). Rituximab elicits its therapeutic effect via direct cell killing, complement activation, and Fc-mediated antibody dependent cellular cytotoxicity (ADCC) (16). In addition, CIT has been previously shown to induce an acute secretory phenotype leading to increased macrophage mediated antibody dependent cellular phagocytosis (ADCP), in a humanized mouse model of B-cell lymphoma (17–19). Therefore, it is important to evaluate the synergistic interaction of BTK inhibition and monoclonal antibody treatment in the context of the tumor microenvironment (TME).

Pre-clinical studies assessing the synergistic interaction between ibrutinib and monoclonal antibodies have provided conflicting results, with a number of studies reporting that ibrutinib not only has no impact on antibody-mediated effects, rather a negative effect specifically on NK-cell mediated effects of the antibody (20–23). Down-regulation of CD20 on CLL cells by ibrutinib has additionally been described in co-culture systems and may be interpreted as a potential antagonistic mechanism (24,25). However, combination therapy of ibrutinib and anti-CD20 antibody has proven to be clinically highly effective especially for unfit patients with CLL (26,27). As the antibody therapy relies on several independent effector mechanisms antibody-dependent cellular phagocytosis has been previously underestimated (28). The effector function of macrophages is highly dependent on interaction within the tumor microenvironment. Therefore, we employed a reliable humanized mouse model of “Double Hit-Lymphoma” for functional elucidation of ibrutinib in antibody combination therapy (17–19).

Here, we show that ibrutinib synergizes with multiple monoclonal antibodies *in vitro* and *in vivo* via increased macrophage mediated ADCP. We screened the kinase activity of malignant B-cells under ibrutinib, tirabrutinib, and acalabrutinib treatment, identifying Janus Kinase 2 (JAK2) as having significantly reduced activity under ibrutinib, whilst second generation BTKis did not similarly inhibit JAK2 activity. Finally, we show using CRISPR/Cas9 knockouts and a kinase inhibitor library that loss of JAK2 as well as inhibition using tofacitinib leads to increased ADCP *in vitro,* whilst superior survival of JAK2 knockout was observed in combination with monoclonal antibody therapy *in vivo.*

## Material and Methods

### Humanized DHL mouse model

For *in vivo* experiments 8-14 weeks old male NOD.Cg-*Prkdc*^scid^ *II2rg*^tm1Wjl^/SzJ (NSG, Jackson Laboratory, USA) immunodeficient mice were injected *i.v.* with 1×10^6^ hMB cells diluted in 100 μl PBS (19). Ten days after injection mice were treated *i.p.* on three consecutive days with ibrutinib (30 mg/kg), alemtuzumab (day 1 1 mg/kg, day 2 and day 3 5 mg/kg), tirabrutinib (30 mg/kg), or PBS as control. Disease progression was monitored by weekly blood sampling and daily scoring of the mice. Spleen and bone marrow were harvested and dissociated by cell strainers with PBS and lysed with 5 ml ACK lysis buffer for 3 min at RT. For flow cytometry samples were stained with F4-80 APC antibody (BioLegend, San Diego, USA).

### Cell lines and primary patient material

The study was approved by the ethical commission of the medical faculty of the University of Cologne (reference no. 13-091) and an informed written consent was obtained from all patients. Primary CLL patient samples were isolated from peripheral blood as previously described (29). To isolate peritoneal macrophages, wild type and BTK^−/−^ C57BL/6 mice were injected *i.p.* with thioglycolate and macrophages obtained via peritoneal lavage after four days(18). BTK^−/−^ C57BL/6 mice were generated by BTK^−/−^ mice backcrossed to C57BL/6J background (30). Peritoneal macrophages and the murine macrophage cell line J774A.1 were cultured in DMEM (Gibco) supplemented with 10 % FBS (Biochrom GmbH) and 1% Pen/Strep (Gibco). The human-MYC/BCL2 (hMB) cell line (strain 102), generated by Leskov et al., was cultured in B-cell culture medium (BCM) composed of a 1:1 ratio of Iscove’s Modified Dulbecco’s Medium (IMDM) and DMEM, supplemented with 10% FBS, 1% P/S, 1% GlutaMAX and 1% β-Mercaptoethanol (19).

### Antibody-Dependent Cellular Phagocytosis (ADCP assay)

J774A.1 macrophages were cultivated in 96-well plates at 1×10^4^ cells per well. After 4 h of incubation at 37°C, 1.5×10^5^ hMB “double-hit” lymphoma cells/well were co-cultured with respective macrophages. Subsequently, this co-culture was treated for 17 h with tyrosine kinase inhibitors and monoclonal antibodies in combination or as mono treatment. Each condition was performed with five replicates. For determination of ADCP GFP^+^ target cells were analyzed using a MACSQuant flow cytometer. The percentage of ADCP was calculated as follows:

100 – (100 x (cells/μL treated / cells/μL untreated)).

### Generation of conditioned media

For the generation of conditioned media 1.5×10^6^ hMB cells/ml were incubated with 0-20 μM of ibrutinib in 10 ml BCM for 24 h. Afterwards the inhibitor was washed off and cells were incubated in 10 ml fresh media for 24 h. Subsequently, the supernatant was taken off for further experiments.

### Phagocytosis F4-80 staining

To determine the amount of phagocyted hMB cells by macrophages, J774A.1 were plated out in a density of 1×10^5^ cells in 1ml media 4 h prior to addition of target cells. Then 1.5×10^6^ hMB cells in 1 ml media, 250 μl alemtuzumab (10 μg/ml) and 250 μl of 0-20 μM ibrutinib were added. After 16 h cells were blocked with 10 μl FcR blocking reagent. Macrophages were stained with F4-80 APC antibody (BioLegend, San Diego, USA) and cells were analyzed using MACSQuant flow cytometry.

### Kinase activity profiling by PamChip microarray

Kinase activity profiles of treated cell lysates were determined via PamChip peptide microarrays on a Pam Station 12 (PamGene International B. V., Netherlands https://www.pamgene.com/en/pamchip.htm). PamChip microarrays contain distinct peptides with 12-15 amino acids representing different proteins. These peptides get phosphorylated depending on the kinase activity in the lysates. Phosphorylated peptides were recognized by phospho-specific PY20 FITC-conjugated antibodies and detected with a CCD camera.

For kinase activity profiling we used Protein-Tyrosine kinase (PTK) and Serine/Threonine (STK) Chips containing 196 and 140 peptides targeting the main kinase families. For PTK activity profiling 5 μg of total protein concentration was loaded on the chip and dissolved in 4 μl 10x PK buffer, 0.4 μl 100x BSA, 4 mM ATP, 0.6 μl FITC conjugated antibody, 10 mM DTT, 4 μl PTK additive and filled up with distilled water to 40 μl total volume (basic mix). All reagents were supplied by PamGene International B.V. For STK activity profiling 1 μg of total protein concentration was loaded on the chip and dissolved in 4 μl 10x PK buffer, 0.4 μl 100x BSA, 4 mM ATP, 0.5 μl STK primary Antibody mix and filled up with distilled water to a total volume of 35 μl (basic mix). 0.4 μl FITC conjugated antibodies dissolved in 3 μl 10x AB buffer and 26.6 μl water were added after an initial incubation time (detection mix). Prior to sample loading a blocking step was performed loading 30 μl of 2% BSA. To determine the kinetic of kinase activity, the samples were pumped several times (cycles) through the microarray and imaged at certain cycle passages. For all experiments biological replicates were used. CLL patient samples were measured in technical replicates.

For data analysis Bio Navigator software (PamGene) was used to calculate the cycle and time dependent signals into a single value for each peptide on the chip (exposure time scaling). Furthermore, outliers due to saturation or insufficient antibody binding were excluded. For analysis, data were log transformed calculating the fold change between treated and untreated, and the p-value was calculated using an unpaired Students’ two-tailed t-test. A p-value of p≤0.05 was accepted as statistically significant. Individual peptides were matched to their representative kinases using a proprietary database (unpublished; PamGene International B. V., Netherlands). In brief, this database ranked the likelihood of peptides belonging to kinases using public databases such as; PhosphoNet, Reactome and UniPROT. Volcano plots were produced using the EnhancedVolcano package in RStudio (R version 3.3.1).

### Statistics

All data was evaluated and graphs generated using GraphPad Prism 8. Unless otherwise stated, bar graphs represent the mean ± SD of three biological replicates. Box plots show the minimal and maximal value, the 25^th^ and 75^th^ quartiles and the median. Statistical comparison between groups was performed using the One-way ANOVA multiple comparison test in the ADCP assays, or the Kruskal-Wallis test for non-gaussian distributed data. Kaplan Meier survival analysis was performed using pairwise Log-rank (Mantel-Cox) test. Differences were considered statistically significant at p-values less than 0.05 (ns=not significant; p > 0.05, p > 0.05; *, p ≤ 0.05; **, p ≤ 0.01; ***, p ≤ 0.001). All statistical analyses were performed using GraphPad Prism version 8.00 (GraphPad Software, San Diego, CA).

## Results

### Ibrutinib enhances macrophage-mediated antibody-dependent cellular phagocytosis

In order to elucidate the potential synergistic interaction between BTKis and monoclonal antibodies on macrophage-mediated ADCP, we utilized multiple monoclonal antibodies (rituximab, obinutuzumab, alemtuzumab), as well as different types of macrophage effector cells in an ADCP co-culture system *ex vivo* with the hMB humanized mouse model of “Double-hit” lymphoma as target cells (17,31) (Figure 1A).

**Figure 1:**
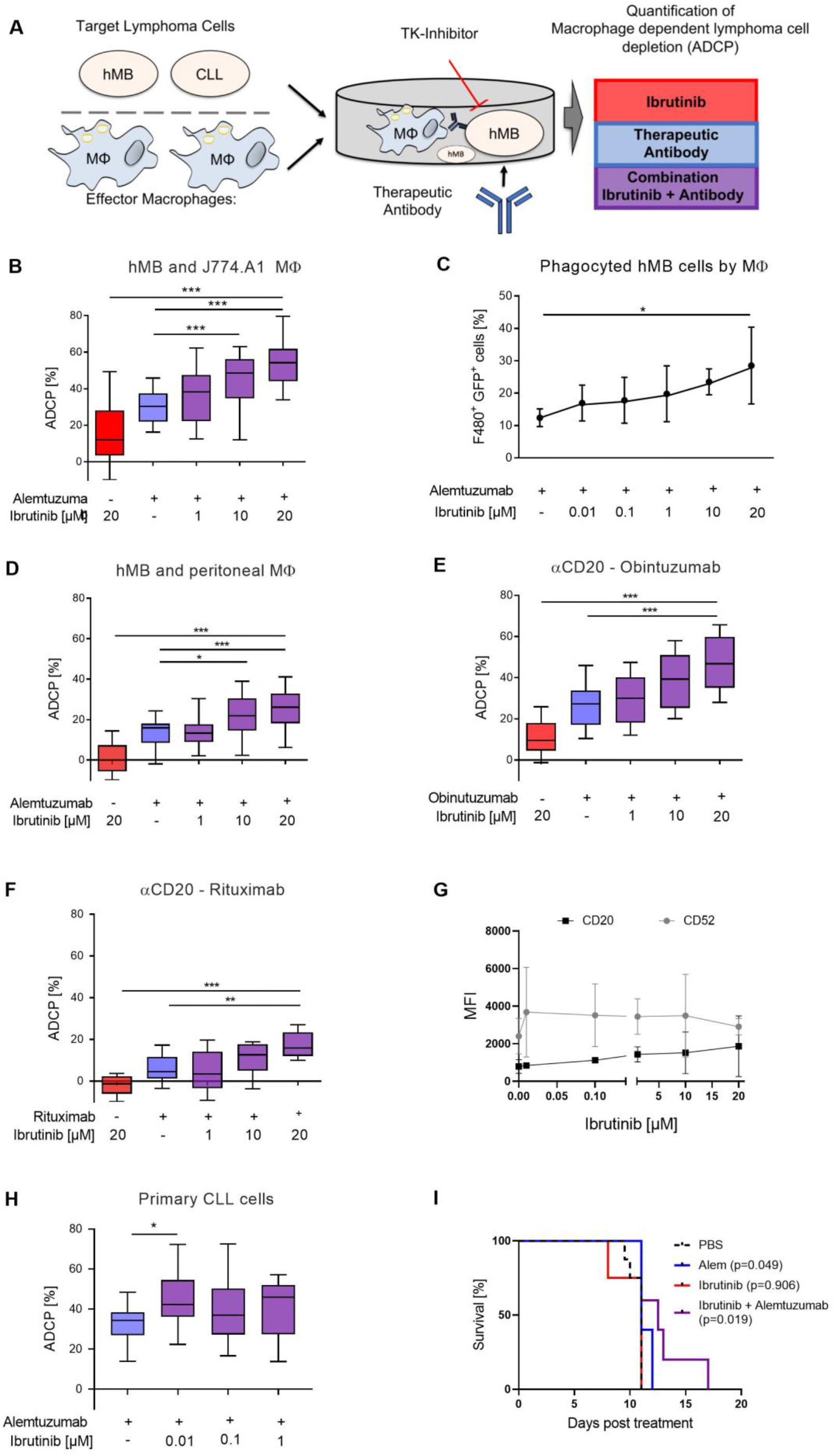
Ibrutinib enhances macrophage-mediated antibody-dependent cellular phagocytosis. **(A)** Schematic of Antibody Dependent Cellular Phagocytosis (ADCP) assay. Cocultures of macrophage effector cells and lymphoma target cells were treated for 17 h with tyrosine kinase (TK) inhibitors and therapeutic antibodies in combination or as monotherapy. Lymphoma target cells were measured by flow cytometry. **(B)** Box plot showing ADCP of hMB “Double-Hit” lymphoma cells and J774A.1 macrophages treated with ibrutinib and alemtuzumab. **(C)** Graph showing phagocyted hMB cells by J774A.1 macrophages, measuring GFP^+^ lymphoma cells and F4/80^+^ macrophages in an ADCP model treated with ibrutinib and alemtuzumab. **(D)** Box plot showing ADCP of hMB lymphoma and murine peritoneal macrophages as effector cells treated with ibrutinib and alemtuzumab. **(E-F)** Box plot showing ADCP of hMB “Double-Hit” lymphoma cells and J774A.1 macrophages treated with ibrutinib and anti-CD20 antibody **(E)** obinutuzumab and **(F)** rituximab. **(G)** Expression of CD20 and CD52 surface markers after ibrutinib treatment of hMB lymphoma cells. **(H)** Box plot showing ADCP of CD19^+^ primary CLL patient cells (N=6) pretreated ex vivo with ibrutinib for 24 h and cocultured with J774A.1 macrophages. alemtuzumab treatment was applied for 24 h. The graphic shows the relative macrophage-dependent cell death in the presence or absence of antibody. **(I)** Kaplan-Meier analysis comparing the survival of male hMB transplanted NSG mice receiving ibrutinib and alemtuzumab as mono therapy or in combination (n≥4 per treatment condition). PBS was used as control. The treatment was given *i.p.* 10 days after *i.v*. hMB cell injection. All box plots show the median, the 25^th^ and 75^th^ quartiles and the minimal and maximal value. All bar graphs display the average and SEM. Unless otherwise stated experiments were performed of at least three biological replicates. (\**P*□<□0.05, \*\**P*□≤□0.01 and \*\*\**P*□≤□0.001)

Co-treatment of alemtuzumab with serially diluted concentrations of ibrutinib lead to significantly increased antibody-mediated lymphoma cell depletion in a concentration dependent manner (Figure 1B). We did not attribute this significant increase to cell toxicity since at these concentrations the cells retained high cell viability (Supp. Figure S1A and B). To verify that ibrutinib specifically enhances phagocytosis of antibody-targeted malignant B-cell lymphoma cells we assessed engulfment of GFP^+^ hMB Double-Hit lymphoma cells into F4/80^+^ J774A.1 macrophages and detected a rising number of F4/80^+^/GFP^+^ cells with increasing ibrutinib concentrations (Figure 1C). Furthermore, this effect was independent of the number of macrophages present under ibrutinib treatment (Supp. Figure S1C). We observed similar significant increases in ADCP performing an independent *ex vivo* experimental setup using primary murine peritoneal macrophages (Figure 1D), whilst also observing concentration dependent antibody-mediated lymphoma cell depletion with the type 1 and 2 anti-CD20 monoclonal antibodies rituximab (Figure 1E) and obinutuzumab (Figure 1F). Thereby, ibrutinib treatment of hMB cells *in vitro* induced a moderate but non-significant increase of CD20 and CD52 expression (Figure 1G). We furthermore evaluated the effects of ibrutinib on primary leukemic cells from chronic lymphocytic leukemia (CLL) patients. Primary CLL cells pretreated with ibrutinib for 24 h exhibited an increased susceptibility towards alemtuzumab mediated ADCP comparing the ibrutinib treatment with antibody to ibrutinib treatment without antibody therapy (Figure 1H). Importantly, here we observe an increase of ADCP using a low ibrutinib concentrations of 0.01 μM. This corresponds to concentrations achieved with oral formulation of 420 mg ibrutinib daily (32).

Finally, we leveraged male NOD.Cg-*Prkdc*^scid^ *II2rg*^tm1Wjl^/SzJ (NSG) mice transplanted with hMB cells and treated them 10 days after transplantation with ibrutinib and alemtuzumab alone and in combination to assess the impact of BTK inhibition on overall survival *in vivo*. Pairwise Kaplan Meier analysis between all treatment groups showed a significant increase in overall survival after alemtuzumab treatment as previously shown (alemtuzumab median = 11 days vs. PBS median = 11 days, p = 0.049) (8) (Figure 1I). Moreover, combination treatment of ibrutinib with alemtuzumab significantly increased overall survival of hMB lymphoma carrying mice (ibrutinib+alemtuzumab median = 12.5 days vs. PBS median = 11 days, p = 0.019) (Figure 1I). In conclusion, we observed that combinatorial treatment of B-cell lymphoma with ibrutinib and monoclonal antibodies increases macrophage-mediated phagocytosis and improves overall survival *in vivo*.

### Ibrutinib elicits increased ADCP independent of BTK inhibition

Previously we have shown that chemotherapy in combination with monoclonal antibody treatment induces an ASAP1 leading to increased macrophage-mediated lymphoma cell depletion *in vivo* (31). Along these lines, we hypothesized that ibrutinib could be eliciting a similar effect. To analyze the impact of soluble factors released by malignant B-cells, conditioned media from ibrutinib pretreated hMB cells was generated and applied to ADCP co-culture of treatment naïve effector and target cells. Here, we observed a significant increase of ADCP induced by conditioned media obtained from ibrutinib pretreated hMB cells (Figure 2A). To validate whether ibrutinib was eliciting its action via the effector or target cells, we conducted pretreatment in both J774A.1 and hMB cells, followed by co-culture with treatment naïve cells in the ADCP assay. When we pretreated macrophages with ibrutinib and subsequently added treatment naïve hMB target cells to the ADCP assay we did not detect any significant change in phagocytosis (Figure 2B). In contrast, when we applied ibrutinib pretreated hMB cells to treatment naïve macrophages in co-culture, we observed a significant increase in ADCP (Figure 2C). This suggests that ibrutinib is eliciting its synergistic interaction with monoclonal antibody mainly in the malignant B-cells.

**Figure 2:**
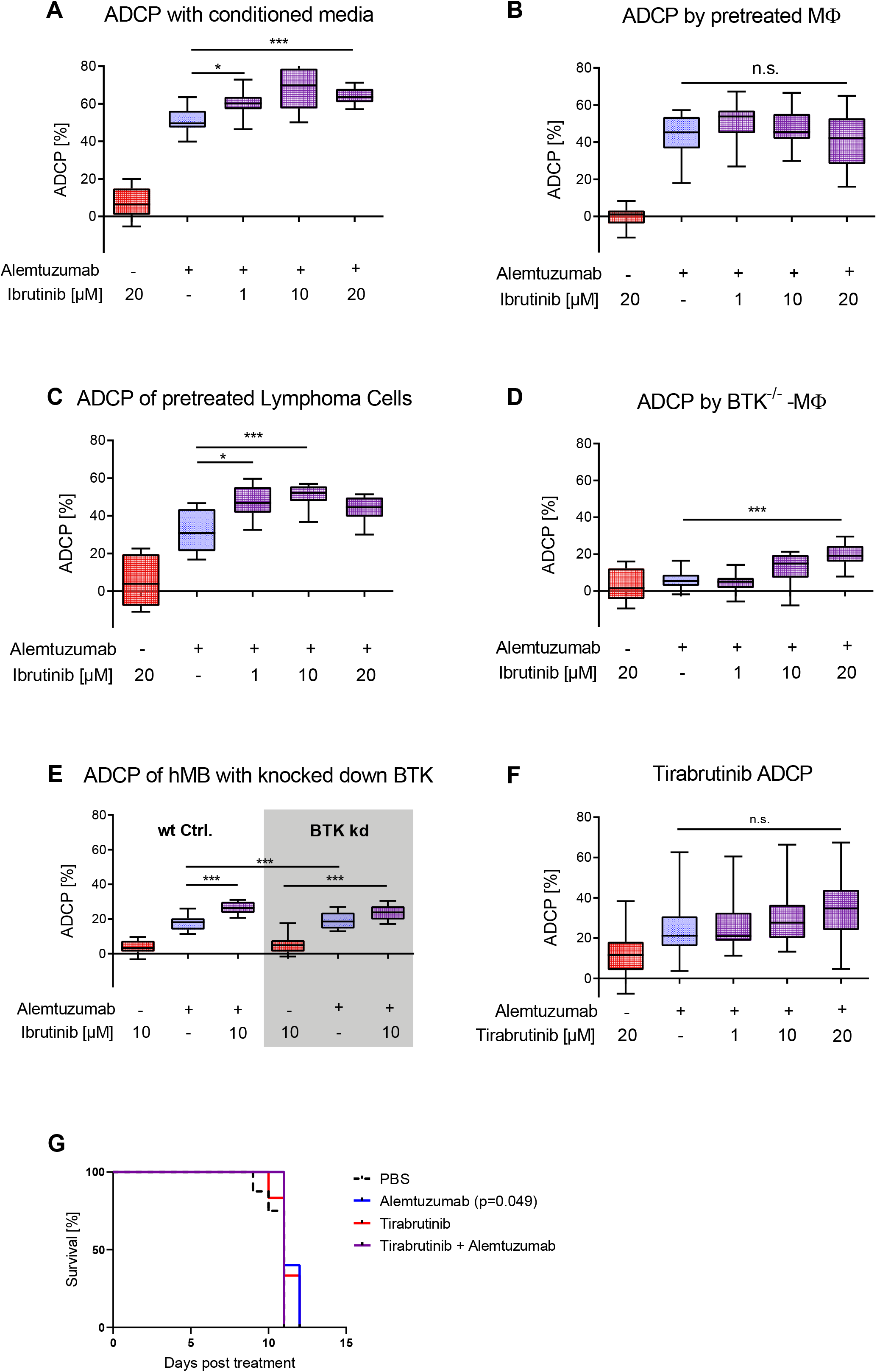
Ibrutinib elicits increased ADCP independent of BTK inhibition. **(A)** Box plot showing ADCP of hMB “Double-Hit” lymphoma target cells and J774A.1 macrophages treated with alemtuzumab and conditioned media of ibrutinib pretreated hMB cells. **(B)** Box plot showing ADCP of hMB lymphoma cells co-cultured with ibrutinib-pretreated J774A.1 macrophages, both treated with alemtuzumab. **(C)** Box plot showing ADCP of ibrutinib-pretreated hMB lymphoma cells co-cultured with J774A.1 macrophages, both treated with alemtuzumab (n=2). **(D)** Box plot showing ADCP of hMB “Double-Hit” lymphoma cells by combination of alemtuzumab with ibrutinib using primary peritoneal macrophages obtained from global BTK^−/−^ mice as effector cells **(E)** Box plot showing ADCP by combination of alemtuzumab with ibrutinib and wildtype (wt) hMB “Double-Hit” lymphoma cells and hMB with knock down in BTK. **(F)** Box plot showing ADCP of hMB “Double-Hit” lymphoma cells and J774A.1 macrophages treated with ibrutinib and second generation BTK inhibitor tirabrutinib. **(G)** Kaplan-Meier analysis comparing the survival of hMB transplanted male NSG mice receiving tirabrutinib and alemtuzumab as mono therapy or in combination (n≥3 per treatment arm). PBS was used as control. The treatment was given i.p. 10 days after i.v. hMB cell injection. All box plots show the median, the 25^th^ 75^th^ quartiles and the minimal and maximal value. Unless otherwise stated experiments were performed of at least three biological replicates. (\**P*□<□0.05, \*\**P*□≤□0.01 and \*\*\**P*□≤□0.001)

In comparison to second generation BTKis, ibrutinib is not as highly selective for BTK binding, having affinity for other Tec kinases, such as EGFR and JAK2 (11). Therefore, we interrogated whether ibrutinib mediated its effect through covalently binding its main target BTK, leveraging primary peritoneal macrophages from BTK^−/−^ mice (Figure 2D) and knocking down BTK in hMB cells (Figure 2E, Supp. Figure S2A). In these experiments we observed significant increases in ADCP for BTK^−/−^ peritoneal macrophages and no significant difference between wt hMB cells and hMB cells with a knock down in BTK treated with alemtuzumab, suggesting that one of ibrutinib’s off-target kinases was responsible for the observed effect. To further confirm that BTK was not responsible for this effect, we conducted ADCP assays *in vitro* using the second generation BTKis tirabrutinib (Figure 2F) and acalabrutinib (Supp. Figure S2B) both showing similar levels of phagocytosis and no toxicity (Supp. Figure S2C and D). Moreover, combination therapy with alemtuzumab and tirabrutinib *in vivo* (Figure 2G) did not increase overall survival and did not reduce hMB cells or macrophages in spleen and bone marrow. (Supp. Figure S2E-H). In conclusion, we have identified that the ADCP-enhancing effects of ibrutinib were mediated by targeting the malignant B-cell compartment, which induces a secretory component in the malignant B-cells leading to macrophage activation. Furthermore, we have shown that the observed effect is independent of BTK, therefore we hypothesized that the increased ADCP induced by ibrutinib to be associated with its off-target kinases.

### Kinase activity profiling identifies Janus Kinase 2 & 3 as the main off-targets for ibrutinib vs. second generation BTKis

To identify the off-target effects of ibrutinib that might be responsible for the synergistic interaction observed in Figures 1 and 2, we analysed the kinase activity profiles of hMB cells as well as CLL patient cells under *in vitro* BTKi treatment after 6 h. Therefore we used PamChip microarrays, an assay which measures the phosphorylation of protein tyrosine as well as serine/threonine kinases (33) (Figure 3A). Treating hMB lymphoma cells with ibrutinib or tirabrutinib predominantly reduced peptide phosphorylation (Figure 3B; left and right hand panels), whilst acalabrutinib-treated cells displayed both up and downregulated peptides (Figure 3B; central panel). We observed 78 significantly altered peptides which were shared across all three BTKis, whereas ibrutinib was the only BTKi to downregulate peptides not observed under the treatment of acalabrutinib or tirabrutinib, indicating the lower specificity of ibrutinib compared to second generation BTKis (Figure 3C). To delve deeper into off-targets of ibrutinib, we conducted a stratified analysis based on kinases that include the same cysteine residue as BTK (34) (Figure 3D). Importantly, all three BTKis targeted BTK in a similar fashion. Furthermore, we observed an enrichment of significant peptides for Janus Kinase (JAK) 2 and 3, as well as for LYN and BLK under ibrutinib treatment, whilst the other Tec kinases showed minimal changes between BTKi treatments. Likewise, ibrutinib treatment of CLL patient cells enrolled significant downregulation of peptides associated with BTK as well as off-target kinases JAK2/3, LYN, BLK and other Tec kinases (Supp. Figure 3A and B). In conclusion, this data suggests that JAK2 and JAK3 could be the off-target kinases of ibrutinib, and therefore potentially the responsible kinases for the increased macrophage-mediated phagocytic capacity.

**Figure 3:**
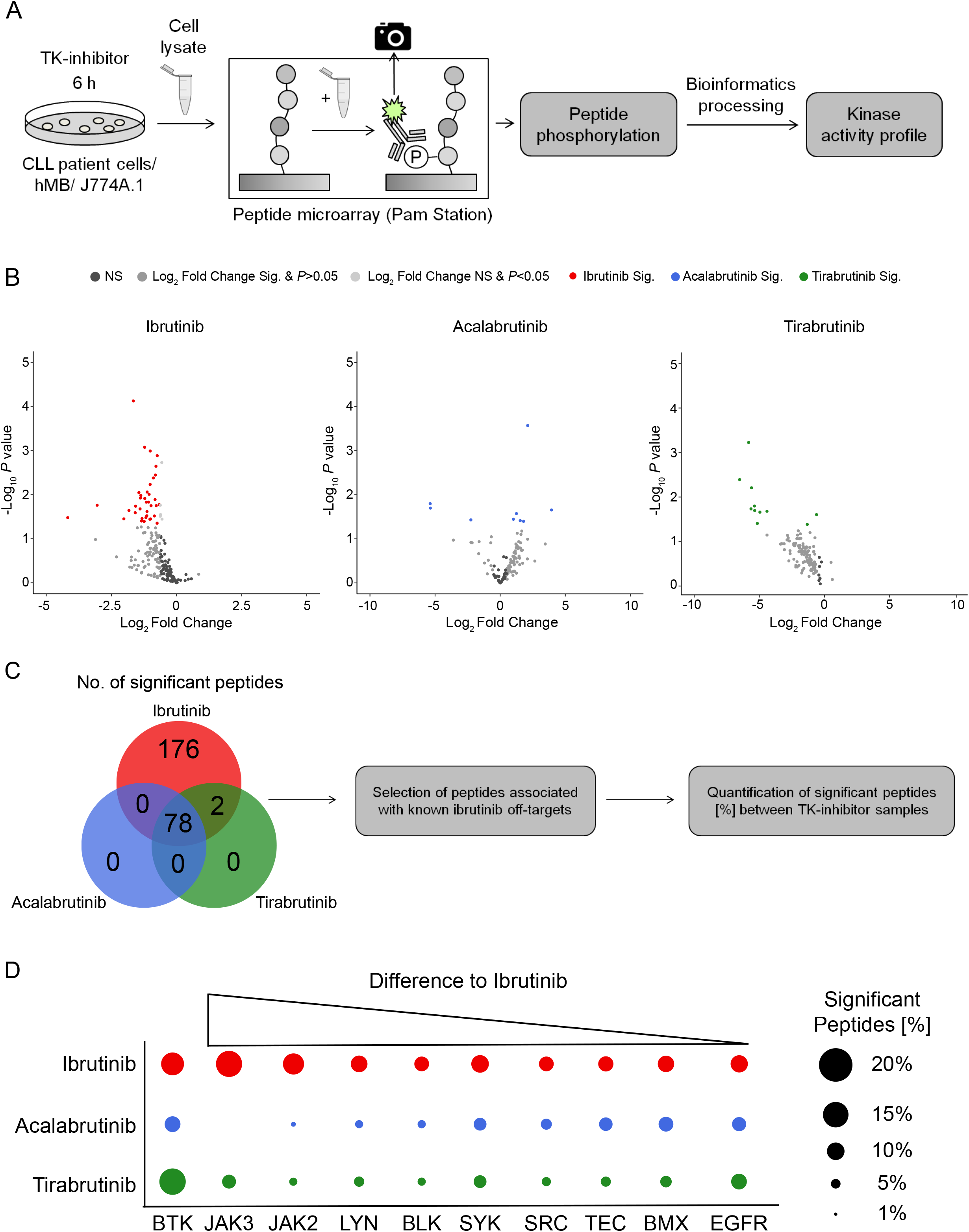
Kinase activity profiling identifies Janus Kinase 2 & 3 as the main off-targets for ibrutinib vs. second generation BTKis. **(A)** Schematic of the PamStation approach assessing lysates of treated cells to peptide microarray chip. Peptides on the chip gets phosphorylated depending on the kinase activity of respective lysates. The phosphorylation is detected via fluorescently labelled antibodies. Bioinformatics processing using different databases transform the peptide phosphorylation level into a kinase activity profile. **(B)** Volcano plots of significantly changed peptide phosphorylation after ibrutinib (red, n=6), acalabrutinib (blue, n=3) and tirabrutinib (green, n=3) treatment of hMB lymphoma cells. Each dot represents a kinase peptide substrate represented on the peptide microarray chip. Coloured dots indicate significantly altered peptides (two-sided students t-test, p ≤ 0.05; log2 fold change ≤ or ≥ 0.5). A negative log2 fold change stands for a downregulation of peptides and a positive log2 fold change for an upregulation compared to the untreated control. Acalabrutinib and tirabrutinib treated lysates were only applied to protein tyrosine kinase (PTK) chips. **(C)** VENN diagram showing all significantly changed peptides of the PTK chips for respective treatments. Based on these significantly changed peptides a kinase activity profile is calculated via bioinformatics processing. **(D)** Graphic showing ibrutinib (red) off-target kinases and its number of significantly changed peptides. Kinase peptide numbers are sorted from the highest difference to acalabrutinib (blue) and tirabrutinib (green) to the lowest. Graphics show biological replicates (n=3).

### JAK2 inhibition with ruxolitinib and tofacitinib enhances macrophage mediated ADCP

To translate our kinase activity findings back to macrophage mediated ADCP, we generated a modestly sized Tec kinase inhibitor library, conducting concentration dependent ADCP assays *in vitro.* As expected, inhibition of EGFR (erlotinib; Figure 4A), SYK (entospletinib; Figure 4B), and BMX (CHMFL-BMX-078; Supp. Figure S4A), lead to no significant increases in phagocytosis. Importantly, inhibition of JAK1/2 via ruxolitinib (Figure 4C), JAK2/3 via tofacitinib (Figure 4D) and pan-JAK inhibition via SP600125 (Supp. Figure S4B) all induced significant increases in macrophage mediated ADCP in a concentration dependent manner. Notably, all inhibitors did not display direct cytotoxicity or apoptosis induction to lymphoma target cells in the concentrations used (Supp. Figure S4C-H).

**Figure 4:**
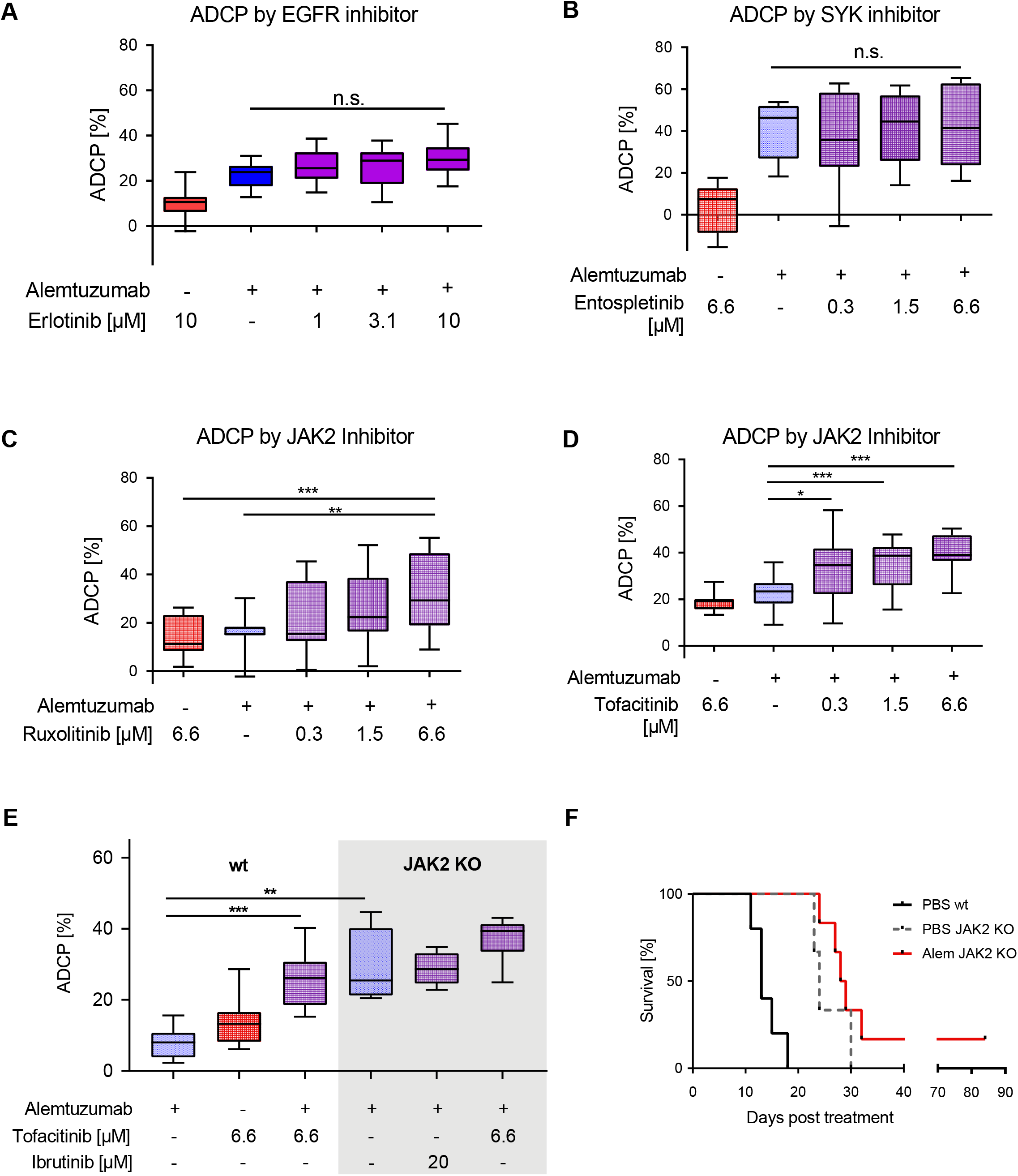
JAK2 inhibition with ruxolitinib and tofacitinib enhances macrophage-mediated ADCP. **(A-D)** Box plot showing ADCP of hMB “Double-Hit” lymphoma cells and J774A.1 macrophages treated with alemtuzumab and **(A)** erlotinib (EGFR inhibitor), **(B)** entospletinib (SYK inhibitor, n=2), **(C)** ruxolitinib (JAK inhibitor) and **(D)** tofacitinib (JAK inhibitor). **(E)** Box plot showing ADCP of JAK2^−/−^ vs empty vector control transduced hMB “Double-Hit” lymphoma target cells and J774A.1 macrophages treated or not treated with JAK-inhibitor tofacitinib, ibrutinib and alemtuzumab (n=2). **(F)** Kaplan-Meier analysis comparing the survival of wildtype (wt) and male JAK2^−/−^ hMB transplanted NSG mice receiving PBS or alemtuzumab (n≥4 per treatment condition). The treatment was given i.p. 10 days after i.v. hMB cell injection (n≥3). All box plots show the median, the 25^th^ and 75^th^ quartiles and the minimal and maximal value. Unless otherwise stated experiments were performed of at least three biological replicates. (\**P*□<□0.05, \*\**P*□≤□0.01 and \*\*\**P*□≤□0.001)

To specifically address the functional implications of JAK2 signaling disruption on lymphoma cell susceptibility to ADCP we generated JAK2 deficient lymphoma target cells (Supp. Figure S4I). In our ADCP assay JAK2^−/−^ cells showed significantly increased phagocytosis under monoclonal antibody treatment in comparison to JAK2 wildtype cells, which could not be improved by co-treatment with either ibrutinib or tofacitinib (Figure 4E). To validate our findings *in vivo,* we leveraged once again hMB humanized mouse model, comparing wildtype hMB cells with JAK2^−/−^ hMB cells with and without alemtuzumab. In this analysis, we observed a clear superior survival of JAK2^−/−^ vs. PBS wildtype cells (PBS JAK2^−/−^ median = 24 days vs. PBS wildtype median = 13 days, p = 0.013) (Figure 4F). Furthermore, alemtuzumab treatment of JAK2^−/−^ cells induced a moderate but not significant increase in survival (alemtuzumab JAK2^−/−^ median = 28,5 days vs. PBS JAK2^−/−^ median = 24 days, p = 0.277). In conclusion, we have shown that inhibition of JAK2 facilitates an increased macrophage-mediated lymphoma cell depletion *in vitro,* with superior overall survival of hMB JAK2^−/−^ transplanted NSG mice *in vivo*.

### Discussion

Her, we have identified a synergistic interaction between ibrutinib and monoclonal antibody treatment on macrophage mediated ADCP in the treatment of B cell lymphoma. Interestingly, we observed that this synergy was not mediated by inhibition of BTK as the primary target of ibrutinib but rather was mediated by targeting of the off-target kinase JAK2. Using JAK2 inhibitors instead of ibrutinib we could recapitulate the synergistic interaction, therefore identifying inhibition of the JAK-STAT signaling pathway as a potential target for antibody combinatorial approaches in the treatment of B cell malignancies.

While monoclonal antibodies exert various mechanisms of targeting malignant cells in B-cell lymphoma, macrophage ADCP has been identified as a pre-dominant mechanism (18,28). In this light, previous reports examining ibrutinib in combination with rituximab that did not reveal synergy were mostly focused on NK-cell ADCC or relied on monocyte derived macrophages (20–23,35). Exploring effector cell mechanisms is technically challenging since co-culture systems demand high levels of standardization. However, regarding NK-cell dependent ADCC only modest effects of ibrutinib have been observed (20) or even debated to be inhibitory (21).

Here we employed two independent macrophage effector cell *in vitro* models including primary peritoneal macrophages. Moreover, our *in vitro* ADCP findings were underlined by significantly improved survival *in vivo* by combination treatment of a humanized mouse model of “Double-Hit” lymphoma (19). As the used NSG mice are immunodeficient and do not display mature T-cells, B-cells or NK cells, the main effector cells are macrophages which mediate the antitumor effect of alemtuzumab and ibrutinib(36).

Off-target effects of ibrutinib have been identified recently by independent methodology (34), likewise using Kinobeads the inhibition of TEC kinase family members and particularly BLK by ibrutinib could be shown (37). However, it remains to be clarified which potential additional kinases beyond BTK are relevant for mediating anti-leukemic effects.

As for the interpretation of clinical trial data it remains to be clarified if ibrutinib combinations with anti-CD20 antibodies rituximab or obinutuzumab are superior to monotherapy in B-cell lymphoma (38). Nevertheless ibrutinib/ rituximab combination therapy has superior outcome to FC-R as the previous first-line standard therapy of CLL (12).

Using second generation BTKis we did not observe a synergistic effect, although the combination of acalabrutinib and obinutuzumab was shown to improve progression-free survival for patients with treatment-naive symptomatic CLL (39). Here, the high single-agent activity of each drug or ADCP-independent mechanisms of synergy are probably reflecting the superior clinical response in CLL. In our work, we primarily address aggressive Double-Hit lymphoma cells with a potentially less prominent dependence of the malignant B-cell towards sustained BTK signaling.

As ibrutinib has shown clinical activity in MCL and non-GCB DLBCL, ibrutinib in combination with R-CHOP regimens for treatment of first-line DLBCL revealed a clinical benefit only in younger patients (40). In Waldenström’s Macroglobulinemia (WM) the combination of rituximab and ibrutinib displayed superior progression-free survival and has been FDA-approved (15). In WM a constitutively activated JAK-STAT signaling pathway has been linked to secretion and disease progression (41). As another B-cell receptor signaling related pathway the impact of JAK2-signaling has also been proposed to be relevant in CLL. Here it could be shown that anti-IgM mediated BCR stimulation induces STAT3 activation signaling in CLL (42). In a clinical trial inhibition of JAK2 by ruxolitinib decreases symptoms (43), however therapeutic impact regarding relevant endpoints such as PFS and OS remains to be clarified. Finally, Kondo et al. could show that ibrutinib treatment of CLL cells also inhibits STAT3 phosphorylation and described suppression of IL10 and PD-L1 in ibrutinib treated CLL patient cells (44). In this line a single cell immune profiling of the temporal dynamics of ibrutinib treatment in CLL patient revealed profound modulation of the non-malignant immune cells and particularly up-regulation of inflammatory pathways in myeloid cells (45). In this context the potential benefits of early phase trials using BTK inhibitors or ruxolitinib in severe COVID-19 patients can be hypothesized to be related to overlapping target specificity and regulation of inflammatory pathways (5,6,46).

In the perspective of B-cell lymphoma we propose JAK2 to be inhibited by ibrutinib and regulating the subsequent phosphorylation of STAT3 and interaction with the tumor microenvironment is particularly leading to re-activation of macrophage effector function. Furthermore, we argue that inhibition of JAK-STAT signaling might sensitize B-cell lymphoma cells towards macrophage mediated ADCP and JAK2 inhibitors should be evaluated and utilized in future treatment concepts of B cell malignancies.

## Acknowledgements

CP was supported by the “NRW -Nachwuchsgruppenprogramm” and Deutsche Forschungsgemeinschaft (KFO286). This work was supported by research funding from Gilead Sciences. We would like to thank Savithri Rangarajan for bioinformatic advice as well as Olaf Merkel and Yvonne Meyer for excellent technical support. Furthermore, we want to thank Max Bondarchenko and Wilfried Ellmeier for providing guide RNAs and BTK^−/^ C57BL/6J mice.

## References

1. Byrd JC, Furman RR, Coutre SE, Flinn IW, Burger JA, Blum KA, et al. Targeting BTK with ibrutinib in relapsed chronic lymphocytic leukemia. N Engl J Med. 2013;

2. O’Brien S, Furman RR, Coutre S, Flinn IW, Burger JA, Blum K, et al. Single-agent ibrutinib in treatment-naïve and relapsed/refractory chronic lymphocytic leukemia: a 5-year experience. Blood. 2018;

3. Wilson WH, Young RM, Schmitz R, Yang Y, Pittaluga S, Wright G, et al. Targeting B cell receptor signaling with ibrutinib in diffuse large B cell lymphoma. Nat Med. 2015;

4. Wang ML, Blum KA, Martin P, Goy A, Auer R, Kahl BS, et al. Long-term follow-up of MCL patients treated with single-agent ibrutinib: Updated safety and efficacy results. Blood. 2015;126:739–45.

5. Treon SP, Castillo J, Skarbnik AP, Soumerai JD, Ghobrial IM, Guerrera ML, et al. The BTK-inhibitor ibrutinib may protect against pulmonary injury in COVID-19 infected patients. Blood. 2020;

6. NCT04346199. Acalabrutinib Study With Best Supportive Care Versus Best Supportive Care in Subjects Hospitalized With COVID-19. CALAVI (Calquence Against the Virus). https://clinicaltrials.gov/show/NCT04346199. 2020;

7. Niemann CU, Wiestner A. B-cell receptor signaling as a driver of lymphoma development and evolution. Semin. Cancer Biol. 2013.

8. Pallasch CP, Hallek M. Incorporating Targeted Agents Into Future Therapy of Chronic Lymphocytic Leukemia. Semin Hematol. 2014;

9. Ponader S, Chen SS, Buggy JJ, Balakrishnan K, Gandhi V, Wierda WG, et al. The Bruton tyrosine kinase inhibitor PCI-32765 thwarts chronic lymphocytic leukemia cell survival and tissue homing in vitro and in vivo. Blood. 2012;

10. Herman SEM, Montraveta A, Niemann CU, Mora-Jensen H, Gulrajani M, Krantz F, et al. The Bruton Tyrosine Kinase (BTK) Inhibitor Acalabrutinib Demonstrates Potent On-Target Effects and Efficacy in Two Mouse Models of Chronic Lymphocytic Leukemia. Clin Cancer Res. 2017;

11. Thompson PA, Burger JA. Bruton’s tyrosine kinase inhibitors: first and second generation agents for patients with Chronic Lymphocytic Leukemia (CLL). Expert Opin. Investig. Drugs. 2018.

12. Shanafelt TD, Wang X V., Kay NE, Hanson CA, O’Brien S, Barrientos J, et al. Ibrutinib-Rituximab or Chemoimmunotherapy for Chronic Lymphocytic Leukemia. N Engl J Med. 2019;381:432–43.

13. Woyach JA, Ruppert AS, Heerema NA, Zhao W, Booth AM, Ding W, et al. Ibrutinib regimens versus chemoimmunotherapy in older patients with untreated CLL. N Engl J Med. 2018;379:2517–28.

14. Burger JA, Sivina M, Jain N, Kim E, Kadia T, Estrov Z, et al. Randomized trial of ibrutinib vs ibrutinib plus rituximab in patients with chronic lymphocytic leukemia. Blood. 2019;133:1011–9.

15. Dimopoulos MA, Tedeschi A, Trotman J, Garcia-Sanz R, Macdonald D, Leblond V, et al. Phase 3 trial of Ibrutinib plus rituximab in Waldenstrom’s macroglobulinemia. N Engl J Med. 2018;

16. VanDerMeid KR, Elliott MR, Baran AM, Barr PM, Chu CC, Zent CS. Cellular cytotoxicity of next-generation CD20 monoclonal antibodies. Cancer Immunol Res. 2018;

17. Lossos C, Liu Y, Kolb KE, Christie AL, van Scoyk A, Prakadan SM, et al. Mechanisms of lymphoma clearance induced by high-dose alkylating agents. Cancer Discov. 2019;

18. Pallasch CP, Leskov I, Braun CJ, Vorholt D, Drake A, Soto-Feliciano YM, et al. Sensitizing protective tumor microenvironments to antibody-mediated therapy. Cell. 2014;156.

19. Leskov I, Pallasch CP, Drake A, Iliopoulou BP, Souza A, Shen C-H, et al. Rapid generation of human B-cell lymphomas via combined expression of Myc and Bcl2 and their use as a preclinical model for biological therapies. Oncogene. 2013;32.

20. Hassenrück F, Knödgen E, Göckeritz E, Midda SH, Vondey V, Neumann L, et al. Sensitive detection of the natural killer cell-mediated cytotoxicity of anti-CD20 antibodies and its impairment by B-cell receptor pathway inhibitors. Biomed Res Int. 2018;

21. Jerkeman M, Ek S, Freiburghaus C, Lindblad AA. Ibrutinib in Combination with Anti-CD20-Antibody Negatively Affects Antibody Dependent Cellular Cytotoxic (ADCC) on Mantle Cell Lymphoma Cell Lines, Not Reversed By the Addition of Lenalidomide. Blood. 2017;

22. Duong MN, Matera EL, Mathé D, Evesque A, Valsesia-Wittmann S, Clémenceau B, et al. Effect of kinase inhibitors on the therapeutic properties of monoclonal antibodies. MAbs. 2015;

23. Da Roit F, Engelberts PJ, Taylor RP, Breij ECW, Gritti G, Rambaldi A, et al. Ibrutinib interferes with the cell-mediated anti-tumor activities of therapeutic CD20 antibodies: Implications for combination therapy. Haematologica. 2015;

24. Pavlasova G, Borsky M, Seda V, Cerna K, Osickova J, Doubek M, et al. Ibrutinib inhibits CD20 upregulation on CLL B cells mediated by the CXCR4/SDF-1 axis. Blood. 2016;

25. Skarzynski M, Niemann CU, Lee YS, Martyr S, Maric I, Salem D, et al. Interactions between ibrutinib and anti-CD20 antibodies: Competing effects on the outcome of combination therapy. Clin Cancer Res. American Association for Cancer Research Inc.; 2016;22:86–95.

26. Francesca Romana Mauro, Stefano Molica, Francesca Paoloni, Gianluigi Reda, Livio Trentin, Monia Marchetti, Daniela Pietrasanta, Marta Coscia, Roberto Marasca, Paolo Sportoletti, Gianluca Gaidano, Lorella Orsucci, Caterina Stelitano, Nicola Di Renzo, Anna RF. IBRUTINIB AND RITUXIMAB AS FRONT-LINE TREATMENT FOR UNFIT PATIENTS WITH CHRONIC LYMPHOCYTIC LEUKEMIA (CLL). PRELIMINARY RESULTS FROM THE GIMEMA LLC1114 STUDY. EHA Libr Mauro F 06/14/19; 266175; PF375.

27. von Tresckow J, Cramer P, Bahlo J, Robrecht S, Langerbeins P, Fink AM, et al. CLL2-BIG: sequential treatment with bendamustine, ibrutinib and obinutuzumab (GA101) in chronic lymphocytic leukemia. Leukemia. 2019;

28. Kurdi AT, Glavey S V., Bezman NA, Jhatakia A, Guerriero JL, Manier S, et al. Antibody-dependent cellular phagocytosis by macrophages is a novel mechanism of action of elotuzumab. Mol Cancer Ther. 2018;

29. Herling CD, Abedpour N, Weiss J, Schmitt A, Jachimowicz RD, Merkel O, et al. Clonal dynamics towards the development of venetoclax resistance in chronic lymphocytic leukemia. Nat Commun. 2018;9.

30. Köprülü AD, Kastner R, Wienerroither S, Lassnig C, Putz EM, Majer O, et al. The Tyrosine Kinase Btk Regulates the Macrophage Response to Listeria monocytogenes Infection. PLoS One. 2013;

31. Pallasch CP, Leskov I, Braun CJ, Vorholt D, Drake A, Soto-Feliciano YM, et al. Sensitizing protective tumor microenvironments to antibody-mediated therapy. Cell. 2014;

32. De Jong J, Sukbuntherng J, Skee D, Murphy J, O’Brien S, Byrd JC, et al. The effect of food on the pharmacokinetics of oral ibrutinib in healthy participants and patients with chronic lymphocytic leukemia. Cancer Chemother Pharmacol. 2015;

33. Kheirollahi V, Wasnick RM, Biasin V, Vazquez-Armendariz AI, Chu X, Moiseenko A, et al. Metformin induces lipogenic differentiation in myofibroblasts to reverse lung fibrosis. Nat Commun. 2019;

34. Berglöf A, Hamasy A, Meinke S, Palma M, Krstic A, Månsson R, et al. Targets for Ibrutinib Beyond B Cell Malignancies. Scand. J. Immunol. 2015.

35. Albertsson-Lindblad A, Freiburghaus C, Jerkeman M, Ek S. Ibrutinib inhibits antibody dependent cellular cytotoxicity induced by rituximab or obinutuzumab in MCL cell lines, not overcome by addition of lenalidomide. Exp Hematol Oncol. BioMed Central Ltd.; 2019;8.

36. Shultz LD, Lyons BL, Burzenski LM, Gott B, Chen X, Chaleff S, et al. Human Lymphoid and Myeloid Cell Development in NOD/LtSz-scid IL2R γ null Mice Engrafted with Mobilized Human Hemopoietic Stem Cells. J Immunol. 2005;

37. Dittus L, Werner T, Muelbaier M, Bantscheff M. Differential Kinobeads Profiling for Target Identification of Irreversible Kinase Inhibitors. ACS Chem Biol. 2017;

38. Robak T, Blonski JZ, Robak P. Antibody therapy alone and in combination with targeted drugs in chronic lymphocytic leukemia. Semin. Oncol. 2016.

39. Sharman JP, Egyed M, Jurczak W, Skarbnik A, Pagel JM, Flinn IW, et al. Acalabrutinib with or without obinutuzumab versus chlorambucil and obinutuzmab for treatment-naive chronic lymphocytic leukaemia (ELEVATE TN): a randomised, controlled, phase 3 trial. Lancet. Elsevier Ltd; 2020;395:1278–91.

40. Younes A, Sehn LH, Johnson P, Zinzani PL, Hong X, Zhu J, et al. Randomized phase III trial of ibrutinib and rituximab plus cyclophosphamide, doxorubicin, vincristine, and prednisone in non–germinal center B-cell diffuse large B-cell lymphoma. J Clin Oncol. 2019;

41. Hodge LS, Ziesmer SC, Yang ZZ, Secreto FJ, Novak AJ, Ansell SM. Constitutive activation of STAT5A and STAT5B regulates IgM secretion in Waldenström’s macroglobulinemia. Blood. 2014;

42. Rozovski U, Wu JY, Harris DM, Liu Z, Li P, Hazan-Halevy I, et al. Stimulation of the B-cell receptor activates the JAK2/STAT3 signaling pathway in chronic lymphocytic leukemia cells. Blood. 2014;

43. Jain P, Keating M, Renner S, Cleeland C, Xuelin H, Gonzalez GN, et al. Ruxolitinib for symptom control in patients with chronic lymphocytic leukaemia: a single-group, phase 2 trial. Lancet Haematol. 2017;

44. Kondo K, Shaim H, Thompson PA, Burger JA, Keating M, Estrov Z, et al. Ibrutinib modulates the immunosuppressive CLL microenvironment through STAT3-mediated suppression of regulatory B-cell function and inhibition of the PD-1/PD-L1 pathway. Leukemia. 2018;

45. Rendeiro AF, Krausgruber T, Fortelny N, Zhao F, Penz T, Farlik M, et al. Chromatin mapping and single-cell immune profiling define the temporal dynamics of ibrutinib response in CLL. Nat Commun. 2020;

46. Cao Y, Wei J, Zou L, Jiang T, Wang G, Chen L, et al. Ruxolitinib in treatment of severe coronavirus disease 2019 (COVID-19): A multicenter, single-blind, randomized controlled trial. J Allergy Clin Immunol [Internet]. 2020; Available from: http://www.sciencedirect.com/science/article/pii/S0091674920307387

