## Supplemental FIgures for "Macrophage-mediated antibody dependent effector function in aggressive B-cell lymphoma treatment is enhanced by Ibrutinib via inhibition of JAK2"

**Supplemental Data**

**Figure S1: Ibrutinib enhances macrophage-mediated antibody-dependent cellular phagocytosis (A)** Toxicity staining of ibrutinib treated hMB lymphoma cells with 7AAD. **(B)** Toxicity staining of ibrutinib treated J774A.1 macrophages with Zombie staining. **(C)** Graph showing F4/80$^+$ J774A.1 macrophages treated with alemtuzumab and different concentrations of ibrutinib. Graphs display the average and SEM. Unless otherwise stated experiments were performed of at least three biological replicates. (**P* < 0.05, ***P* ≤ 0.01 and ****P* ≤ 0.001)

**Figure S1**

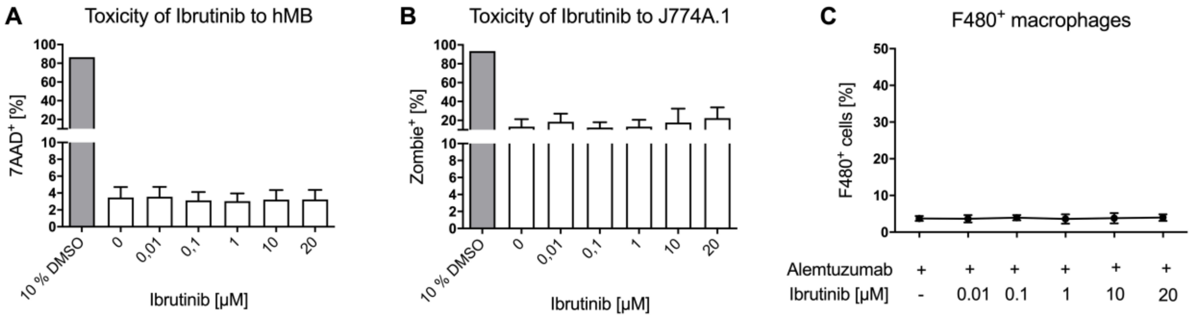

Figure S2

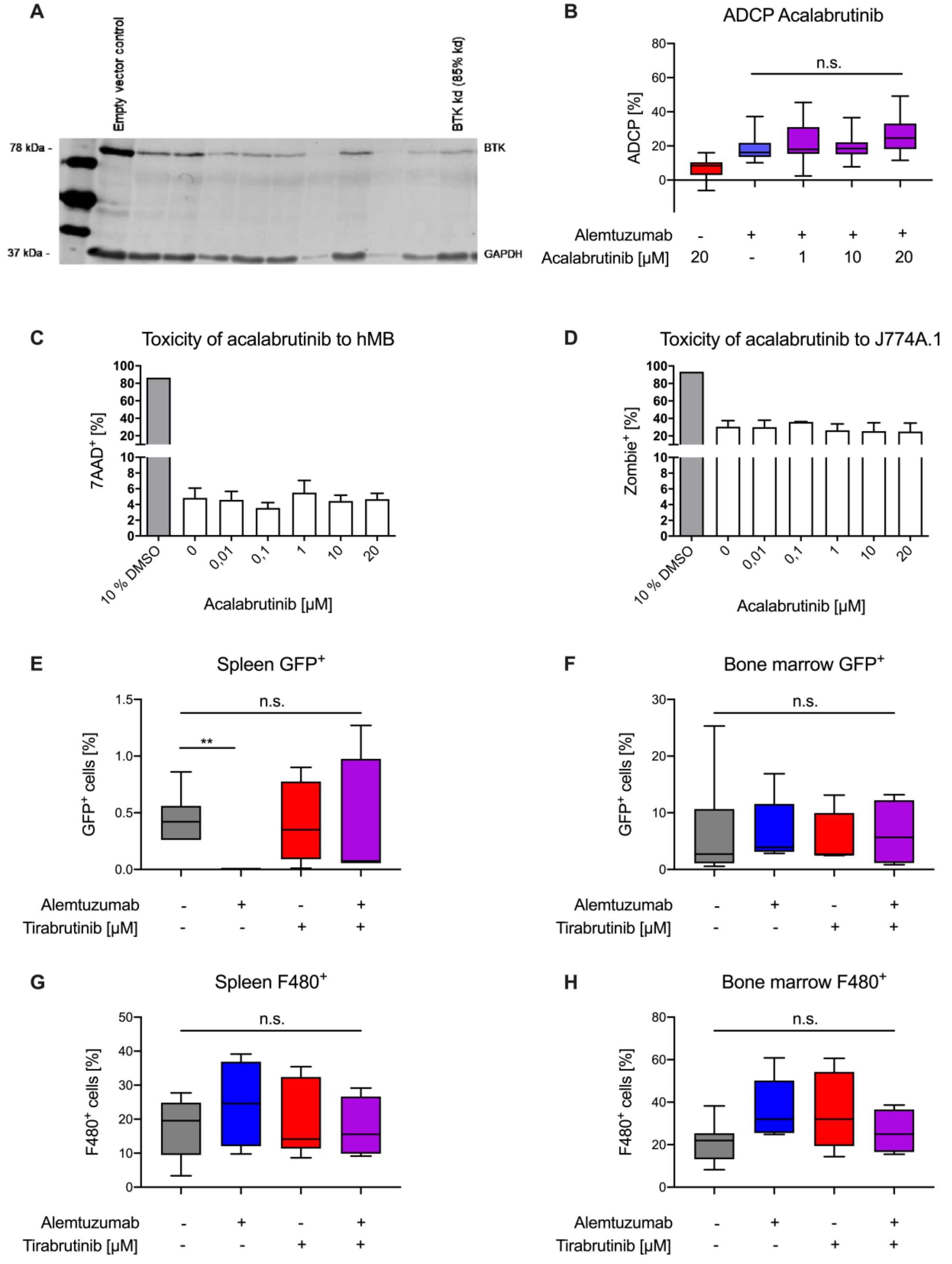


**Figure S2: Ibrutinib elicits increased ADCP independent of BTK inhibition (A)**

Western blot analysis of BTK expression in mCHERRY+-sorted control vector-infected versus BTK vector-infected cells. GAPDH serves as loading control. hMB “Double-Hit” lymphoma cells showed a knock down of 85 %. **(B)** Box plot showing ADCP of hMB lymphoma cells and J774A.1 macrophages treated with alemtuzumab and acalabrutinib (2$^{nd}$ generation BTKi). **(C)** Toxicity staining of acalabrutinib treated hMB lymphoma cells with 7AAD. **(D)** Toxicity staining of acalabrutinib treated J774A.1 macrophages with Zombie staining. **(E-F)** Box plot showing GFP$^{+}$ hMB lymphoma cells in the **(E)** spleen and **(F)** bone marrow after survival of male hMB transplanted NSG mice treated with alemtuzumab and tirabrutinib in combination or as monotherapy. **(G-H)** Box plot showing F4/80$^{+}$ macrophages in the **(G)** spleen and **(H)** bone marrow of male hMB transplanted NSG mice with respective treatments. All box plots show the median, the 25$^{th}$ and 75$^{th}$ quartiles and the minimal and maximal value. All bar graphs display the average and SEM. Unless otherwise stated experiments were performed of at least three biological replicates. (*$P < 0.05$, **$P \leq 0.01$ and ***$P \leq 0.001$)

Figure S3

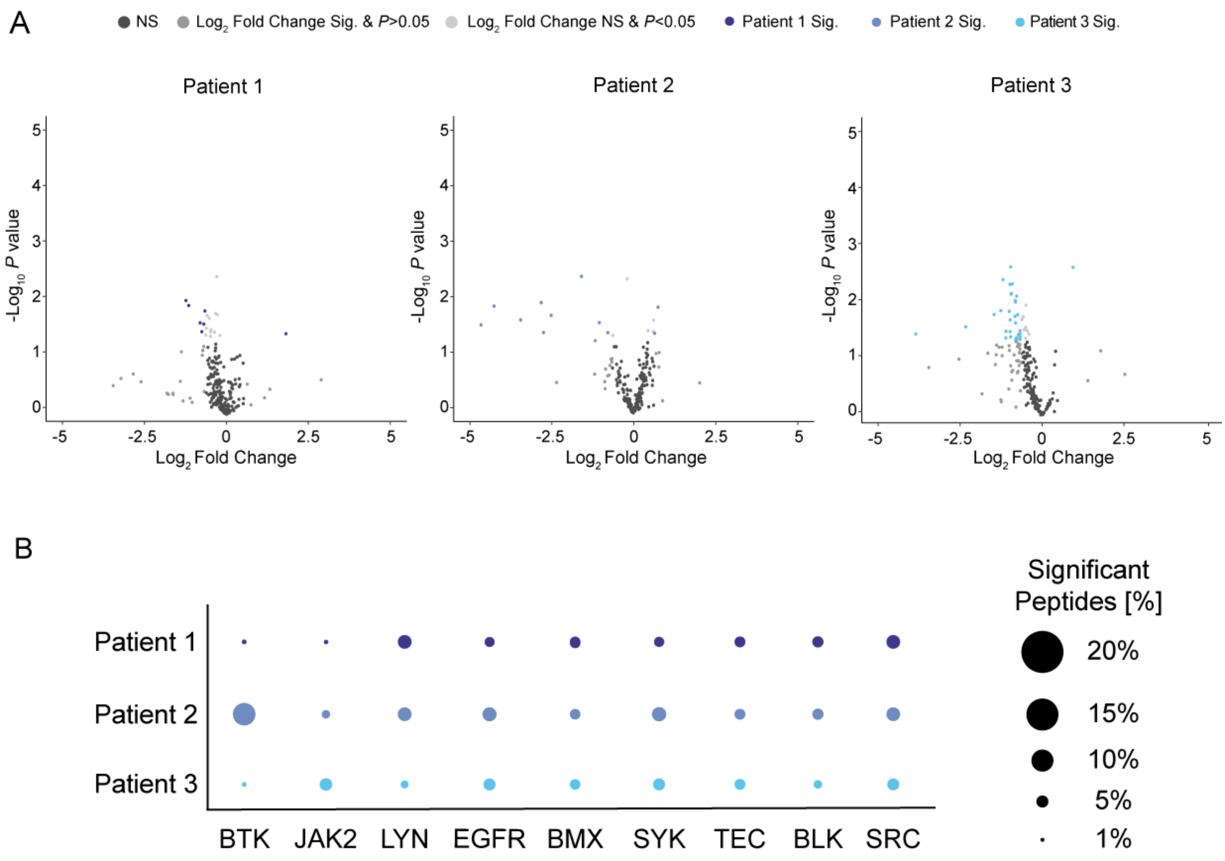


**Figure S3: Kinase activity profiling of CLL patient cells identifying the main off-targets for ibrutinib (A)** Volcano plot of significantly changed peptide phosphorylation after ibrutinib treatment of CLL patient cells. Each dot represents a kinase peptide substrate on the peptide microarray chip. Coloured dots indicate significantly altered peptides (two-sided students t-test, $p \leq 0.05$; log2 fold change ≤ or ≥ 0.5). A negative log2 fold change stands for a downregulation of peptides and a positive log2 fold change for an upregulation compared to the untreated control. **(B)** Graphic showing ibrutinib off-target kinases and its number of significantly changed peptides. Graphics show technical replicates (n=3).

**Figure S4**

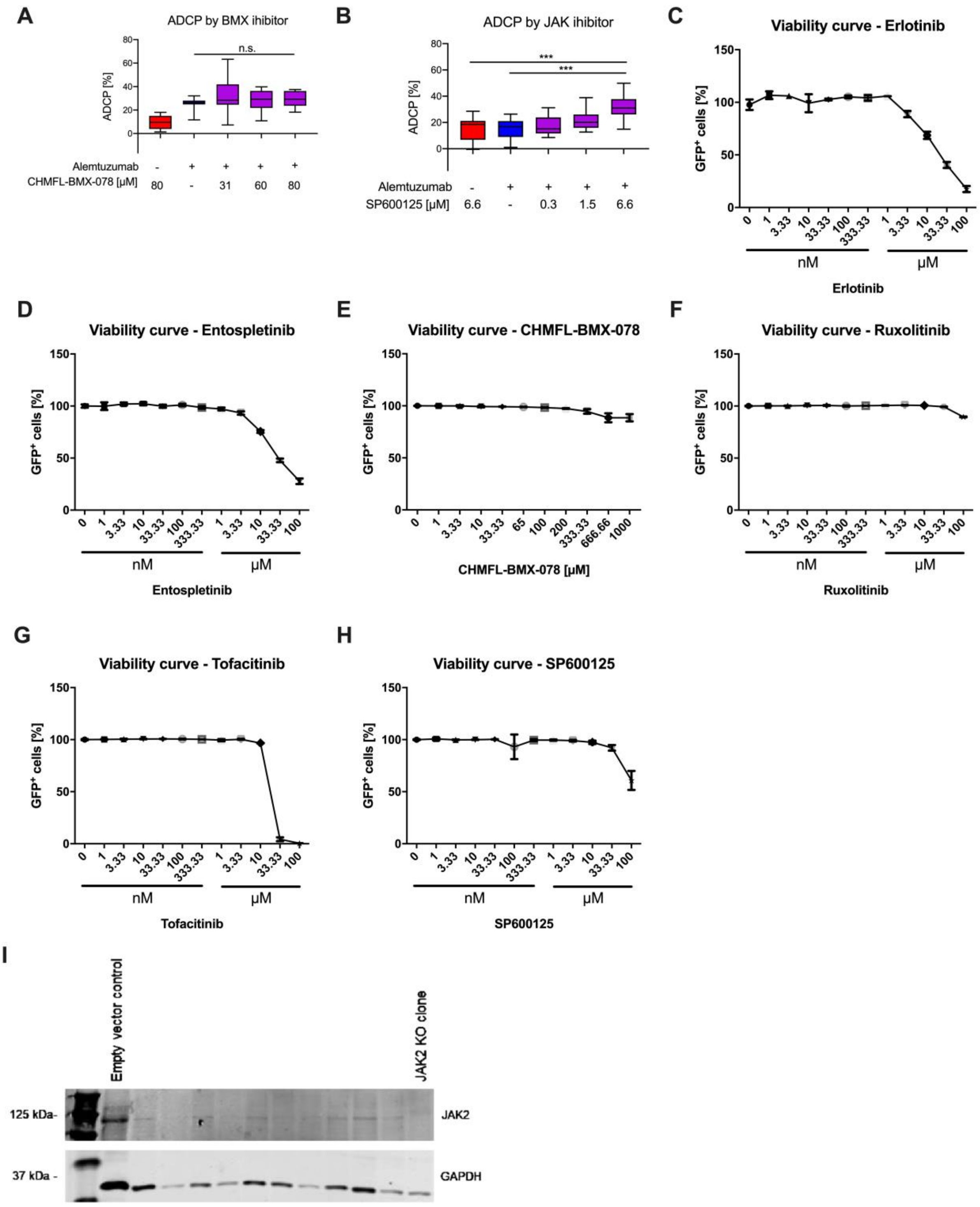


**Figure S4: JAK2 inhibition with ruxolitinib and tofacitinib enhances macrophage-mediated ADCP (A-B)** Box plot showing ADCP of hMB “Double-Hit” lymphoma cells and J774A.1 macrophages treated with alemtuzumab and **(A)** CHMFL-BMX-078 (BMX inhibitor, n=2) or **(B)** SP600125 (JAK inhibitor). **(C-H)** Viability curve of GFP$^+$ hMB lymphoma cells treated with **(C)** erlotinib (EGFR inhibitor, n=2), **(D)** entospletinib (SYK inhibitor), **(E)** CHMFL-BMX-078 (BMX inhibitor), **(F)** ruxolitinib (JAK inhibitor), **(G)** tofacitinib (JAK inhibitor) and **(H)** SP600125 (JAK inhibitor). **(I)** Western blot analysis of JAK expression in mCHERRY$^+$-sorted control vector-infected versus JAK2$^{-/-}$-infected cells. GAPDH serves as loading control. All box plots show the median, the 25$^{th}$ and 75$^{th}$ quartiles and the minimal and maximal value. Viability curves show the mean and SEM. Unless otherwise stated experiments were performed of at least three biological replicates. (*$P < 0.05$, **$P \leq 0.01$ and ***$P \leq 0.001$)

### Supplemental Methods

#### Cell lysate preparation and protein quantification

For lysate preparation 5x10$^6$ hMB, J774A.1 macrophages and CLL patient cells were treated for 6 h with either 1 µM ibrutinib, acalabrutinib or tirabrutinib. Afterwards cells were washed with PBS and centrifuged for 5 min at 300xg. Next 100 µl M-PER lysis buffer (Mammalian Extraction Buffer, ThermoFisher Scientifc #78503, #78420, #87785) with 1 µl Halt Phosphatase Inhibitor Cocktail (100x, Thermo Fischer Scientific #78428) and 1 µl Halt Protease Inhibitor Cocktail, EDTA free (100x, Thermo Fischer Scientific #78437) was given to the cell pellet and incubated for 15 min on ice. Then the cell pellets were centrifuged for 15 min at 16.000×g at 4 °C. The supernatant containing the proteins was transferred into a fresh tube and frozen at -80 °C in 10 µl aliquots. Protein concentration was determined using Pierce BCA Protein Assay (Thermo Fisher, #23225) and measured with microplate reader FluoStar Optima (BMG Labtech).

#### Toxicity staining

To analyze toxicity, 1x10$^6$ cells in 2 ml medium were incubated with respective tyrosine kinase inhibitor for 24 h. After 24 h, macrophages were stained with Zombie NIR$^M$Fixable Viability

Staining diluted 1:100 in PBS and incubated for 15 min in the dark at RT. hMB and CLL patient cells were stained with 7AAD diluted 1:100 in PBS for 15 min at 4°C. 7-AAD toxicity staining was measured by MACSQuant flow cytometer.

### Generation of CRISPR mediated knock-out in hMB cells

All guide RNAs for BTK knock out (KO) (#7707 1-4), JAK2 KO (#7572 6-7) and non-target plasmids (#80248) were gifted from John Doench and David Root[1]. First, hMB lymphoma cells were transduced with a virus containing CRISPR mCherry Cas9 guide RNA gifted from Agata Smogorzewska (#99154) and afterwards with the respective KO guide RNA. For Virus production amphotrophic HEK 293T phoenix cells were used. The hMB cell line was lentivirally transduced by spin infection. 1x$10^6$ cells/ml were plated out on a 12 well plate in 1 ml media. Afterwards, 1ml of respective virus containing medium was added, together with polybrene (2µg/ml final concentration) and centrifuged for 45min at 400g. Then, cells were cultured and antibiotic selection with 10 µg/ml puromycin was started for at least 1 week. After a week cell sorting and western blot analysis was performed.
